# Human brains implicitly and rapidly distinguish AI from human voices before decoding prosodic meaning

**DOI:** 10.64898/2026.04.08.716483

**Authors:** Wenjun Chen, Marc D. Pell, Xiaoming Jiang

**Affiliations:** Institute of Language Sciences, Shanghai International Studies University, Shanghai, 201620, China; Key Laboratory of Language Science and Multilingual Artificial Intelligence, Shanghai International Studies University, Shanghai, 201620, China; School of Communication Sciences and Disorders, McGill University, Montréal, H3G 1A8, Canada

**Keywords:** AI voices, prosody, person perception, EEG, neural decoding, representational similarity analysis

## Abstract

People encounter AI voices daily. Existing behavioral studies suggest listeners rely on prosodic cues such as intonation and expressiveness to detect audio deepfakes, reporting that AI voices sound prosodically less rich than human voices. To test whether prosodic processing drives deepfake discrimination in the neural time course of voice processing, we recorded electroencephalographic (EEG) data while participants listened to human and AI-generated speakers producing utterances in confident vs. doubtful prosody (tone of voice), with attention directed toward memorizing speaker names. We used voice cloning to control for speaker identity confounds between human and AI voices. Multivariate pattern analysis revealed that neural discrimination of human vs. AI voices emerged rapidly regardless of prosody (confident: 176 ms; doubtful: 134 ms), substantially preceding prosody discrimination (confident vs. doubtful within human voices: 2066 ms; within AI voices: 1366 ms). Acoustic analysis confirmed that prosodic distinctions became classifiable only at utterance offset (90% normalized duration), converging with neural evidence that prosody requires near-complete temporal integration. This temporal dissociation between rapid voice source discrimination and late-emerging prosody decoding suggests that prosody plays a smaller role in audio deepfake detection than listeners retrospectively report.

Representational similarity analysis further revealed that spectral envelope features (mel-frequency cepstral coefficients; MFCC), rather than the visually salient high-frequency energy differences, drove neural human–AI discrimination, with MFCC’s earliest independent contribution (228 ms) closely following the MVPA decoding onset (134–176 ms). Future studies may manipulate specific acoustic components to establish the causal sources of this rapid and sustained neural discrimination.

**Significance Statement:** People encounter AI voices daily, in phone calls, navigation apps, supermarket checkouts, and subway announcements. Using electroencephalography, we show that the human brain automatically and rapidly distinguishes everyday AI voices from human speech, even without conscious attention to voice source. Although people may attribute this ability to AI voices sounding monotone or prosodically unnatural, the brain relies on subtler acoustic signatures, enabling discrimination before prosodic information becomes available. Attempts to identify the specific acoustic features driving this neural detection were inconclusive, pointing to the need for future causal investigations. We encourage engineers and policymakers to ensure AI voices remain perceptually detectable, as increasingly humanlike AI voices could cognitively disadvantage the general public if they become indistinguishable from human speech.

## 1. Introduction

When receiving a phone call that starts with “*Hello, I received your resume*,” some people might immediately recognize it as an audio deepfake because the intonation of “*Hello*” already sounds unnatural. This aligns with findings from the literature showing that listeners rely heavily on prosodic cues (e.g., intonation, stress, rhythm) when asked how they distinguish human from AI-generated speech, as human voices are typically perceived as more expressive (Kühne et al., 2020; Rodero, 2017; San Segundo et al., 2025). Possibly due to an incomplete understanding of prosodic cues’ contribution to audio deepfake detection, a recent large-scale video deepfake study manipulated only the visual modality (Groh et al., 2022), despite audio being an essential component of video. Against this background, the present study investigates whether human vs. AI voice detection depends on prosodic discrimination.

We first consider findings in prosody prototypes. Using reverse-correlation techniques in which listeners judged voices with randomly perturbed fundamental frequency (F0) contours, prior work found that dominance judgments rely on lower overall pitch combined with gradual pitch decrease (Ponsot et al., 2018a), while certainty and honesty share a common rising-falling prosodic signature (Goupil et al., 2021). However, these findings were based on single words, and whether such prototypes generalize to sentence-level speech has not been established (Knight et al., 2018). Directly comparing F0 contours across prosody types is more challenging in utterances than in single words due to variability in length and linguistic content, so instead we collected indirect evidence by examining when prosodic distinctions become decodable acoustically and neurally.

To test whether the brain discriminates human from AI voices, and if so, when and whether this discrimination depends on prosodic cues, we followed the established framework for person perception from speech (Lavan et al., 2024) and collected EEG data while participants listened to human vs. AI-generated speech (Roswandowitz et al., 2024) in confident vs. doubtful prosodies (Jiang et al., 2020). Voice cloning was used to match AI voices to each human speaker’s identity, controlling for speaker-specific acoustic confounds that would otherwise prevent attribution of neural discrimination to voice source per se (Chen et al., 2025a, 2026; Roswandowitz et al., 2024). Critically, listeners were instructed to memorize each voice’s name (Lavan et al., 2019) and ignore other stimulus features, ensuring voice source and prosody perception remained task-irrelevant. Implicit paradigms of this kind are necessary to dissociate automatic neural discrimination from conscious detection strategies, as explicit deepfake detection tasks may instead reflect top-down decision-making (Beauchemin et al., 2006; Ma et al., 2023; Pinheiro et al., 2023; Rinke et al., 2022).

Temporal multivariate pattern analysis (MVPA) (Hausfeld et al., 2012; Li Calzi et al., 2025; Sarrett & Toscano, 2024; van de Nieuwenhuijzen et al., 2016) was applied to decode voice source (human vs. AI) and prosody (confident vs. doubtful) from EEG activity at each timepoint, enabling direct comparison of when each dimension becomes neurally discriminable. To further identify the acoustic basis of neural human–AI discrimination, we conducted representational similarity analysis (RSA) (Giordano et al., 2021; Jiang & Pell, 2024; Khalighinejad et al., 2017; Kriegeskorte et al., 2008; Lavan et al., 2024) comparing neural representational geometry with acoustic feature models (F0, high-frequency energy, and spectral envelope features).

We hypothesized that if prosody drives audio deepfake detection, voice source discrimination should be absent or emerge no earlier than prosody decoding; if behavioral reports instead reflect retrospective attribution, voice source decoding should substantially precede prosody decoding. As the acoustic signatures distinguishing human from AI voices remain incompletely characterized (Bisogni et al., 2024), RSA was used to explore their acoustic basis.

## 2. Materials and methods

### 2.1. Participants

Forty native Mandarin speakers (20 females, age: 22.4 ± 1.7 years; 20 males, age: 23.2 ± 1.7 years) were recruited. None reported speech, hearing, or neurological/psychiatric disorders. All provided written informed consent. This study was conducted in accordance with the Declaration of Helsinki. The ethical committee of the Institute of Language Sciences, Shanghai International Studies University, approved the experiment (20230628027). Participants were compensated at 50 RMB/hour.

### 2.2. Stimuli

The 192 training stimuli were drawn from a validated corpus of 11,808 Mandarin Chinese audio recordings (Chen et al., 2026). Twenty-four native Mandarin speakers (12 females, 12 males) were recorded in a sound-attenuated laboratory (Audio-Technica AT2035 microphone; Komplete Audio 6 Mk2 sound card) producing 15 sentences under neutral, confident, and doubtful prosody conditions, following established elicitation procedures (Jiang & Pell, 2017). These recordings were used to train speaker- and style-specific AI voice avatars via Huawei’s Celia system, with each prosodic condition cloned independently (Chen & Jiang, 2023). The trained AI models then generated readings of 123 novel Mandarin sentences expressing factual information, evaluations. Approximately one month later, the original 24 speakers returned to the laboratory under identical recording conditions to produce matched human versions of the same 123 sentences, having heard the AI-generated versions as prosodic references. All files were normalized to −30 dBFS (44,100 Hz). Independent perceptual validation (N = 48 raters) confirmed that human voices were rated as substantially more humanlike than AI voices, and confident prosody received higher confidence ratings than doubtful prosody for both human and AI voices (Chen et al., 2026). For the present study, 192 training stimuli were selected from the 960-stimulus EEG set (Chen et al., 2025b), with speakers organized into four height-matched groups to control for height-related acoustic cues, and stimuli selected to ensure perceptual discriminability across voice source and prosody conditions.

### 2.3. Experimental Design

The experiment comprised eight blocks (human: blocks 1–4; AI: blocks 5–8), with speaker gender further blocked within each source half (male speakers: blocks 1–2, 5–6; female speakers: blocks 3–4, 7–8). Each block contained three phases: Training, Checking, and Testing (Chen et al., 2025b). The full experiment lasted 90–100 minutes including breaks. During Training, participants learned to associate three speakers’ voices with Chinese surnames at their own pace, hearing 12 unique sentences twice each (24 trials per block; 192 trials total across 8 blocks). Each participant heard 96 unique utterances comprising 48 human voice utterances (blocks 1–4) and their perfectly matched AI counterparts (blocks 5–8; identical speaker, sentence, and prosody condition). To counterbalance prosody-stimulus associations, two stimulus sets were created (participants 1–20: Set 1; participants 21–40: Set 2), with the prosody condition of every speaker reversed between sets. The present study analyzes EEG data from the Training phase only. Checking phase accuracy is also reported to confirm attentiveness to speaker–name associations. See **Figure 1A**.

**Figure 1.**
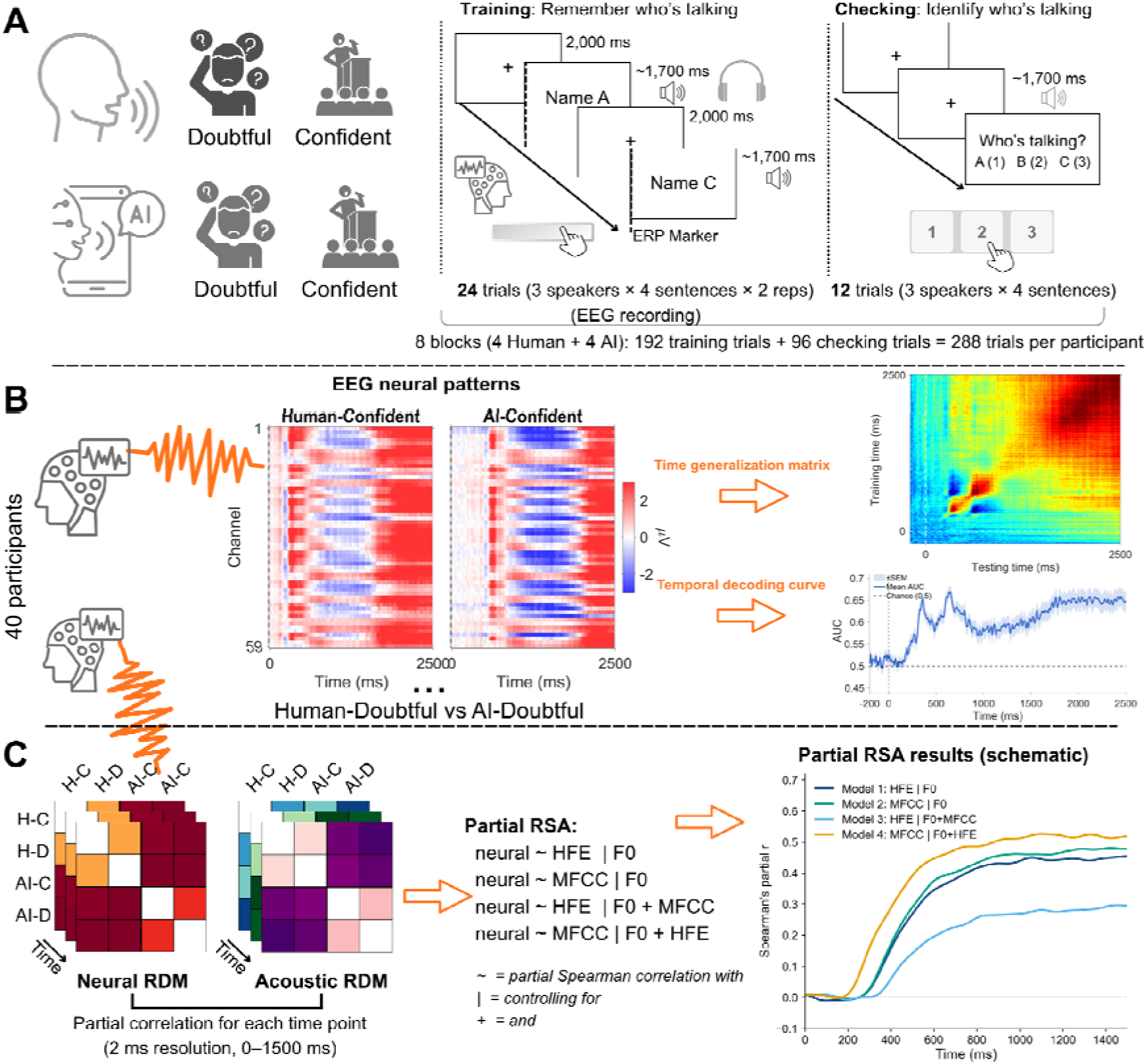
Experimental Design and Analysis Pipeline. **(A)** Speaker identity learning task across 8 blocks (4 Human, 4 AI), each comprising a Training phase (24 trials; EEG recorded) and a Checking phase (12 trials). **(B)** MVPA was applied to EEG patterns (59 channels) from the Training phase to decode voice source (Human vs. AI) at each timepoint using linear discriminant analysis, yielding temporal decoding curves (AUC) and time generalization matrices. **(C)** Partial RSA compared neural RDMs (4×4, correlation distance; 2 ms resolution, 0–1500 ms) with acoustic feature RDMs (F0, HFE, MFCC). Four models assessed independent contributions of HFE and MFCC controlling for F0 and each other (M1: HFE|F0; M2: MFCC|F0; M3: HFE|F0+MFCC; M4: MFCC|F0+HFE). Schematic time courses illustrate the expected pattern of results.

### 2.4. EEG Recording

Continuous EEG was recorded in a sound-treated, dimly lit, electromagnetically shielded room using a 64-channel elastic cap with passive Ag/AgCl electrodes (ActiCap, Brain Products, Germany) arranged according to the extended 10–20 system, referenced online to FCz. Electrode impedances were maintained below 5 kΩ using conductive gel. Signals were digitized at 500 Hz (bandpass: 0.01–100 Hz). Two EOG electrodes placed near the left outer canthus and below the right eye monitored horizontal and vertical eye movements and blinks. Auditory stimuli were delivered binaurally via insert earphones. Participants sat approximately 80 cm from a computer monitor and were instructed to minimize movement and blink only between trials. A video demonstration of the experimental setup and recording procedure is available (Chen & Jiang, 2024).

### 2.5. Statistical Analyses

#### 2.5.1. Acoustic and Perceptual Analyses of The Stimuli

To verify that the 192 stimuli maintained acoustic comparability and perceptual discriminability across conditions, we first analyzed mean F0 and perceptual ratings of perceived confidence and humanlikeness. Since mean F0 is a primary acoustic marker of speaker identity (Kinoshita et al., 2009; Lavan et al., 2019), it was extracted from vowel-only sequences of each audio file using the extractvowels plugin from Praat’s Vocal Toolkit (Corretgé, 2024) in Praat 6.2.09 (Boersma & Weenink, 2021). Mean F0 was analyzed using linear mixed-effects models (LMER) in R (version 4.3.3) with the lmerTest package (Kuznetsova et al., 2017): `Mean_F0 ∼ Source × Prosody × Gender + (1|Speaker) + (1|Item)`. For perceptual validation, independent raters evaluated all 192 stimuli on 7-point Likert scales for perceived confidence (1 = not confident at all, 7 = very confident) and perceived humanlikeness (1 = very machine-like, 7 = very humanlike), using models: `Perceived_confidence ∼ Source × Prosody + (1|Speaker) + (1|Item)` and `Perceived_humanlikeness ∼ Source × Prosody + (1|Speaker) + (1|Item)`. Simple effects were examined using the emmeans package (Lenth et al., 2021) and effect sizes were quantified using Cohen’s *d*.

To examine when prosodic distinctions become acoustically decodable across utterance duration, time-resolved machine learning classification was applied to F0 contours extracted from all 192 training stimuli (**Figure 2C**). F0 was extracted using *librosa*’s YIN algorithm (McFee et al., 2025) after removing leading and trailing silences. Each F0 contour was time-normalized to 100 equally-spaced points (0–100%), z-score normalized, and smoothed using a Savitzky-Golay filter (window = 15, polynomial order = 3). Random Forest classifiers (100 trees, maximum depth = 5; scikit-learn) (Pedregosa et al., 2011) were trained to distinguish confident from doubtful prosody separately for human and AI voices at 5% intervals from 5% to 100% of normalized utterance duration, using a 10% temporal window centered at each time point. Performance was evaluated using 5-fold cross-validation and assessed against chance level (50%) using one-sample *t*-tests with false discovery rate (FDR) correction for multiple comparisons (Benjamini & Hochberg, 1995).

**Figure 2.**
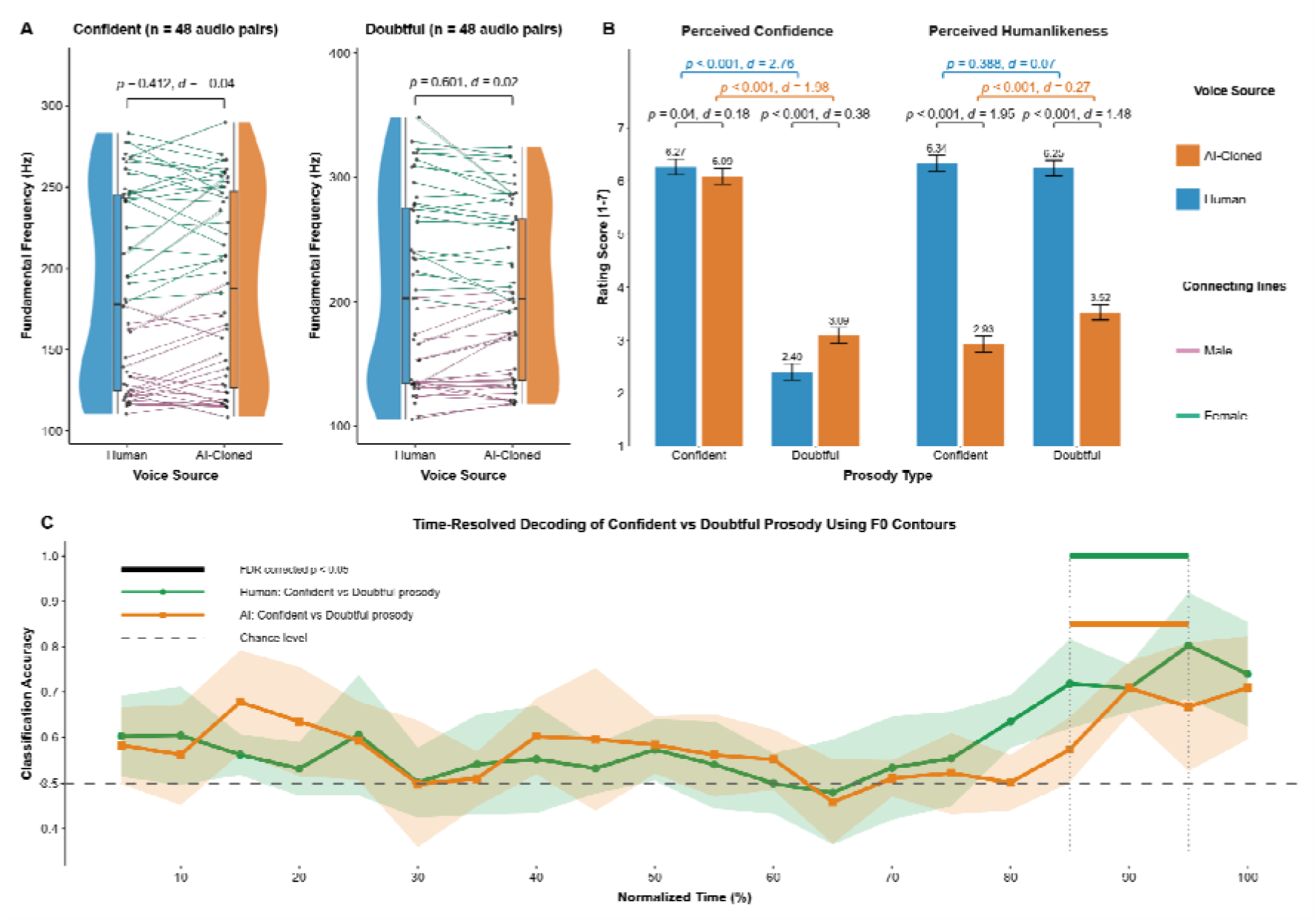
Validation of training stimuli. **(A)** Mean F0 comparison between AI and human voices in two prosodies. Connecting lines link matched audio pairs (same prosody, statement, speaker). **(B)** Perceptual ratings of humanlikeness and vocal confidence (1-7 scale; error bars: SE). **(C)** Machine learning classification of confident vs. doubtful prosody at 5% intervals (5-100% normalized duration). Filled markers indicate FDR-corrected significance (*p* < 0.05).

To illustrate the acoustic properties of representative stimuli, mel spectrograms were computed for one matched human-AI speaker pair across both prosodic conditions using the *librosa* package (version 0.11.0) (McFee et al., 2025) in Python. Spectrograms were computed with a sampling rate of 22,050 Hz, an FFT window size of 2,048 samples (Hamming window), a hop length of 512 samples, and 128 mel frequency bands. Power spectrograms were converted to decibel scale relative to peak amplitude. To enable direct visual comparison of acoustic patterns across utterances, leading and trailing silences were standardized to 50 ms for all displayed utterances using *pydub* (Robert, 2021).

#### 2.5.2. EEG Data Preprocessing and Multivariate Pattern Analysis

Preprocessing was conducted using EEGLAB 2025.0 (Delorme & Makeig, 2004) in MATLAB R2024b. Continuous EEG was re-referenced offline from FCz to the average of bilateral mastoid electrodes (TP9/TP10). To ensure precise time-locking to acoustic onset, leading silence durations were quantified for each audio file using *pydub* (Robert, 2021) in Python 3.11.7, and EEG event markers were adjusted accordingly to account for acoustic onset differences between human and AI recordings. Data were bandpass filtered at 0.1–40 Hz. Bad channels and trials were identified and rejected through visual inspection. Independent component analysis (ICA) was performed on 1 Hz high-pass filtered data; the resulting weights were transferred to the 0.1 Hz filtered data for artifact removal. ICA components reflecting eye movements, muscle activity, and other non-neural artifacts were identified using ICLabel (Pion-Tonachini et al., 2019) and removed following manual inspection. Data were subsequently low-pass filtered at 30 Hz. The online reference electrode (FCz), bilateral mastoid reference electrodes (TP9/TP10), and two EOG channels (FT9/FT10) were removed, yielding 59 channels for analysis. Epochs were defined from −500 to 2598 ms relative to acoustic onset and baseline-corrected using the −500 to 0 ms pre-stimulus interval.

After artifact rejection, a mean of 186.1 trials per participant were retained (*SD* = 13.4, range: 127–192, out of 192 maximum). Trial retention was balanced across conditions: Human-Confident (46.3 ± 4.2), Human-Doubtful (46.7 ± 2.8), AI-Confident (46.1 ± 5.1), and AI-Doubtful (47.0 ± 2.7). Across all 40 participants, 7,443 trials were included in the final analysis (Human-Confident: 1,851; Human-Doubtful: 1,866; AI-Confident: 1,845; AI-Doubtful: 1,881).

Time-resolved MVPA was conducted using the MVPA-Light toolbox (Treder, 2020) in MATLAB R2024b. Two decoding targets were examined: voice source (human vs. AI) and prosodic category (confident vs. doubtful). For voice source decoding, classifiers were trained separately within each prosody condition (confident and doubtful); for prosody decoding, classifiers were trained separately within each voice source (human and AI).

Temporal decoding was performed using linear discriminant analysis (LDA) via `mv_classify_across_time`, with 5-fold cross-validation repeated 5 times. Performance was quantified using area under the receiver operating characteristic curve (AUC), with chance level at 0.5. Statistical significance was assessed using cluster-based permutation testing (Maris & Oostenveld, 2007) with 1,000 permutations and one-sample t-tests against chance, controlling family-wise error rate at α = .05. The cluster-forming threshold was set at |t| = 1.96.

To identify the acoustic basis of neural voice source discrimination, time-resolved representational similarity analysis (RSA) (Kriegeskorte et al., 2008) was conducted over 0–1500 ms post-stimulus onset at 2 ms resolution. ***Neural RDMs.*** For each participant, EEG epochs were averaged within each of four conditions (Human-Confident, Human-Doubtful, AI-Confident, AI-Doubtful) at each timepoint, yielding condition-averaged spatial patterns (59 channels). Pairwise correlation distances between the four condition patterns were computed at each timepoint, resulting in a 4×4 neural representational dissimilarity matrix (RDM) per timepoint per participant. Individual RDMs were averaged across participants to yield a group-mean neural RDM. ***Acoustic Feature RDMs.*** Three acoustic features were extracted from all 192 training stimuli using a 50 ms sliding window interpolated onto the EEG time axis (500 Hz): fundamental frequency (F0), extracted using *librosa*’s (McFee et al., 2025) YIN algorithm (75–500 Hz); high-frequency energy (HFE, >4 kHz), computed as mean spectral power above 4 kHz; and mel-frequency cepstral coefficients (MFCC, 13 coefficients) (Davis & Mermelstein, 1980). Features were averaged within each of the four conditions, and pairwise Euclidean distances between condition-averaged vectors were computed at each timepoint, yielding three 4×4 acoustic RDMs (F0, HFE, MFCC). ***Model RDMs.*** Two binary model RDMs were constructed: a source RDM (human vs. AI; off-diagonal cells = 1 for cross-source pairs, 0 otherwise) and a prosody RDM (confident vs. doubtful; constructed analogously) (Nili et al., 2014).

##### Partial RSA

To assess the degree to which acoustic features predict neural voice source structure independent of prosodic structure, partial Spearman correlations were computed per participant per timepoint between the neural RDM and each acoustic feature RDM, controlling for the remaining predictors (Lavan et al., 2024). Source RSA correlated each acoustic RDM with the source model RDM controlling for the prosody model RDM; Prosody RSA tested the reverse. Partial correlations were computed using the residual method (rank ➔ regress ➔ correlate). To further dissociate the independent contributions of HFE and MFCC beyond F0, four additional partial RSA models were specified: Model 1 (HFE | F0), Model 2 (MFCC | F0), Model 3 (HFE | F0 + MFCC), and Model 4 (MFCC | F0 + HFE). Group-level significance was assessed using one-sided sign-flip cluster permutation testing (10,000 permutations, Z > 1.645, α = .05). Between-model comparisons used two-sided testing (Z > 1.96).

### 2.6. Data and Code Availability

Data and code are available at https://osf.io/chje7/overview?view_only=ed137660b3e74abab321ee214a323c35.

## 3. Results

### 3.1. Listeners Were Focused on Speaker Identity During EEG Recording

We examined recognition accuracy in the Checking phase, where participants identified speakers using a three-alternative forced-choice task (12 trials per block). Overall accuracy was 94.2% (±3.1% SD across subjects), significantly above chance level (33.3%), confirming active engagement in identity learning. Accuracy varied by condition: Human-Confident (95.2% ± 5.6%), Human-Doubtful (95.2% ± 5.1%), AI-Confident (91.6% ± 5.0%), and AI-Doubtful (94.7% ± 5.3%). Notably, human voices showed higher accuracy than AI voices, particularly in the confident prosody condition, resonating with our human-first block ordering to establish a baseline for comparison.

High Checking phase accuracy confirms that listeners successfully directed their attention toward speaker identity learning during the Training phase, as intended.

### 3.2. Comparable Mean F0, Perceptually Distinguishable Ratings, and Late-Emerging Prosodic Classification from F0 Trajectories

We first confirmed that human and AI voices were acoustically and perceptually comparable (**Figure 2A-B**). F0 did not differ significantly between AI and human versions under either confident (*p* = 0.412, *d* = -0.04) or doubtful (*p* = 0.601, *d* = 0.02) prosody. Perceptual ratings confirmed successful prosodic manipulation: both human (*p* < 0.001, *d* = 2.76) and AI (*p* < 0.001, *d* = 1.98) speakers’ confident utterances were rated significantly higher in perceived confidence than doubtful utterances. Human voices were rated significantly more humanlike than AI voices in both confident (*p* < 0.001, *d* = 1.95) and doubtful (*p* < 0.001, *d* = 1.48) conditions.

Then, machine learning classification of F0 contours (**Figure 2C**) revealed that prosodic discriminability emerged only at 90% of normalized duration for both human (70.8%) and AI voices (70.9%), indicating that prosodic information becomes decodable near utterance offset.

### 3.3. Voice Source Was Decoded Within Milliseconds While Prosodic Discrimination Emerged Only Near Utterance Offset

We then investigated neural time courses for discriminating voice source and prosody. We presented mel spectrograms of representative audio samples in **Figure 3A-D**. Visually, human voices (**A**, **C**) exhibited more diffuse high-frequency energy (> 4 kHz), whereas AI voices (**B**, **D**) showed reduced high-frequency components with smoother spectral patterns. Neurologically, our MVPA TGM in **Figure 3E-H** visually revealed that classifiers trained on late-phase neural activity generalized better to late-phase testing time points for both human vs. AI discrimination and prosody discrimination, although this late-phase generalization was stronger for human vs. AI discrimination.

**Figure 3.**
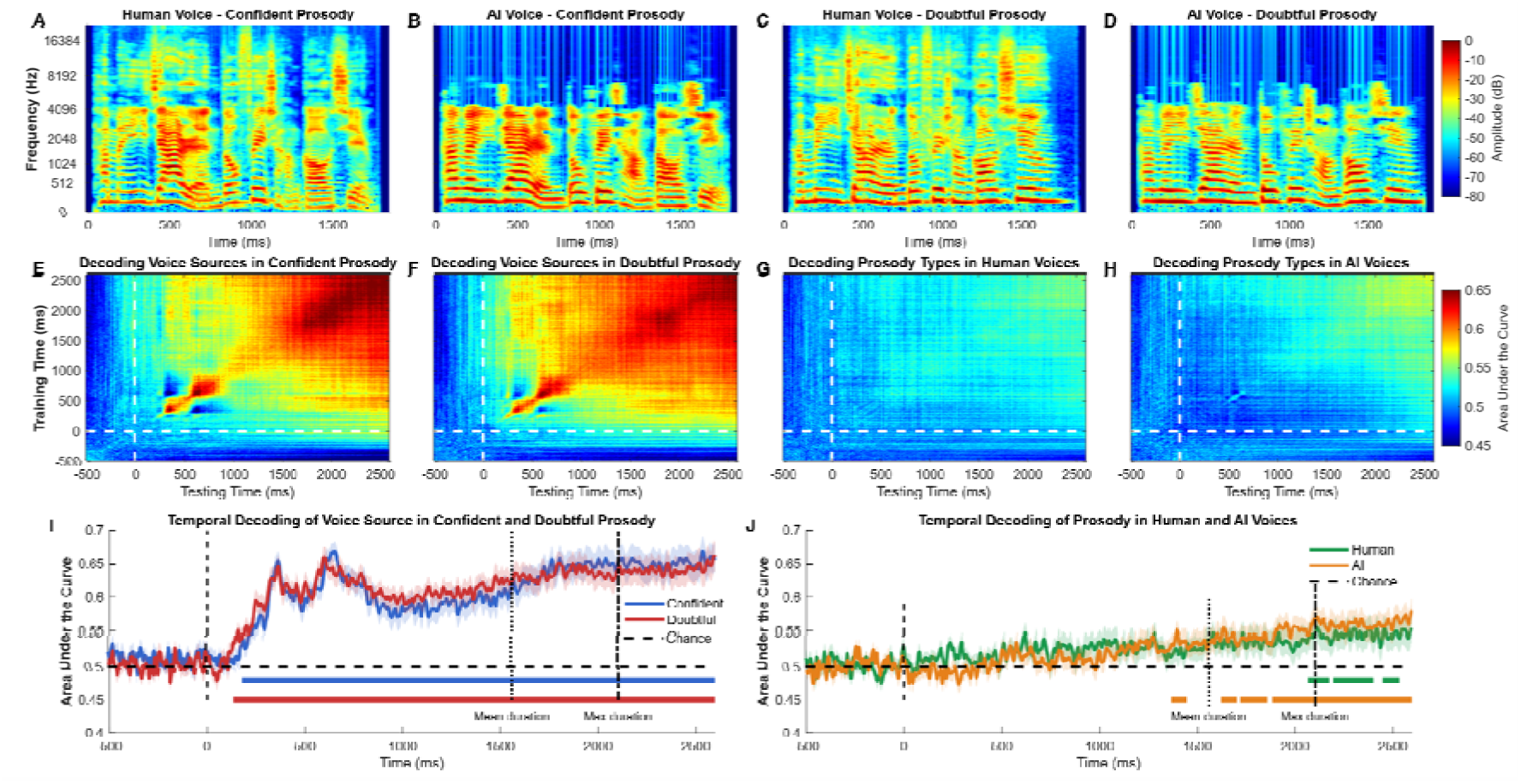
Mel spectrograms and temporal decoding of voice source and prosody. **(A-D)** Spectrograms of an exemplar sentence across conditions. **(E-H)** Time-generalization matrices (TGM) showing decoding performance. **(I-J)** Temporal decoding curves. Shaded areas: SE. Colored bars: significant clusters (*p* < 0.05, cluster-based permutation). Dashed lines: mean (1559 ms) and maximum (2103 ms) stimulus duration.

Statistically, temporal decoding curves (**Figure 3I-J**) showed that, first, listeners’ brain activity significantly decoded human vs. AI voice source under both confident (onset: 176 ms, cluster: 176-2598 ms, *p* < 0.001, peak AUC = 0.669 at 652 ms) and doubtful (onset: 134 ms, cluster: 134-2598 ms, *p* < 0.001, peak AUC = 0.662 at 2594 ms) prosody conditions. Second, prosody decoding emerged substantially later in both voice sources. Within human voices, discrimination onset occurred at 2066 ms, with three significant clusters (2066-2176 ms, *p* = 0.035; 2194-2400 ms, *p* = 0.024; 2448-2532 ms, *p* = 0.039; peak AUC = 0.556 at 2112 ms). Within AI voices, discrimination began earlier at 1366 ms, with four significant clusters (1366-1444 ms, *p* = 0.047; 1622-1702 ms, *p* = 0.046; 1718-1858 ms, *p* = 0.020; 1882-2598 ms, *p* = 0.001; peak AUC = 0.579 at 2596 ms).

We annotated stimulus durations in **Figure 3I-J** and found that (1) human vs. AI discrimination emerged rapidly (134-176 ms) in both prosodic conditions; (2) within human voices, prosody decoding onset (2066 ms) occurred near the maximum duration (2103 ms); and (3) within AI voices, prosody decoding began earlier (1366 ms), closer to the mean duration (1559 ms).

### 3.4. Despite Visually Salient High-Frequency Differences, Spectral Envelope Features Drove Neural Human–AI Voice Discrimination

As illustrated in **Figure 3A–D**, human voices exhibited more diffuse high-frequency energy (>4 kHz) compared to AI voices, which showed reduced and smoother high-frequency spectral patterns. This motivated a systematic examination of whether HFE and MFCC independently drive neural human–AI discrimination beyond F0.

F0, HFE, and MFCC all independently predicted neural voice source structure when controlling for prosody structure (**Table 1**). F0 showed significant clusters from 312 ms (5 clusters, largest: 452–1086 ms, *p* < .001, peak *r* = 0.64 at 642 ms). HFE emerged earlier at 234 ms (7 clusters, largest: 518–786 ms, *p* < .001, peak *r* = 0.52 at 734 ms), and MFCC similarly from 238 ms (7 clusters, largest: 578–702 ms, *p* < .001, peak *r* = 0.51 at 636 ms). Notably, no acoustic feature predicted neural prosody structure (Prosody RSA: all *p* > .05).

**Table 1.** Source and Prosody RSA: Neural Representational Structure Predicted by Acoustic Features.

| RSA type | Feature | Cluster | Onset (ms) | Offset (ms) | Duration (ms) | $p$ | Peak $r$ | Peak time (ms) |
| --- | --- | --- | --- | --- | --- | --- | --- | --- |
| Source RSA | F0 | 1 | 312 | 364 | 52 | 0.02 | 0.49 | 358 |
| Source RSA | F0 | 2 | 378 | 448 | 70 | 0.01 | 0.4 | 390 |
| Source RSA | F0 | 3 | 452 | 1086 | 634 | < .001 | 0.64 | 642 |
| Source RSA | F0 | 4 | 1092 | 1226 | 134 | 0.01 | 0.3 | 1110 |
| Source RSA | F0 | 5 | 1280 | 1500 | 220 | < .001 | 0.42 | 1454 |
| Source RSA | HFE | 1 | 234 | 288 | 54 | 0.01 | 0.45 | 260 |
| Source RSA | HFE | 2 | 388 | 428 | 40 | 0.01 | 0.38 | 412 |
| Source RSA | HFE | 3 | 436 | 508 | 72 | 0.01 | 0.37 | 458 |
| Source RSA | HFE | 4 | 518 | 786 | 268 | < .001 | 0.52 | 734 |
| Source RSA | HFE | 5 | 792 | 908 | 116 | < .001 | 0.49 | 812 |
| Source RSA | HFE | 6 | 1234 | 1360 | 126 | < .001 | 0.39 | 1320 |
| Source RSA | HFE | 7 | 1382 | 1484 | 102 | 0.00 | 0.3 | 1478 |
| Source RSA | MFCC | 1 | 238 | 294 | 56 | 0.01 | 0.36 | 260 |
| Source RSA | MFCC | 2 | 342 | 390 | 48 | 0.01 | 0.37 | 374 |
| Source RSA | MFCC | 3 | 408 | 444 | 36 | 0.02 | 0.38 | 412 |
| Source RSA | MFCC | 4 | 578 | 702 | 124 | < .001 | 0.51 | 636 |
| Source RSA | MFCC | 5 | 710 | 770 | 60 | 0.00 | 0.46 | 748 |
| Source RSA | MFCC | 6 | 1000 | 1056 | 56 | 0.02 | 0.23 | 1026 |
| Source RSA | MFCC | 7 | 1254 | 1376 | 122 | < .001 | 0.3 | 1326 |
| Prosody RSA | F0 | - | - | - | - | > .05 | - | - |
| Prosody RSA | HFE | - | - | - | - | > .05 | - | - |
| Prosody RSA | MFCC | - | - | - | - | > .05 | - | - |

*Note.* One-sided cluster permutation tests (10,000 permutations, $Z > 1.645$ , $\alpha = .05$ ). Source RSA controls for prosody structure; Prosody RSA controls for source structure. No significant clusters for Prosody RSA (0–1500 ms). HFE = high-frequency energy (>4 kHz); MFCC = mel-frequency cepstral coefficients (13 coefficients).

To dissociate the independent contributions of HFE and MFCC beyond F0, we examined Models 1–4 (**Figure 4** , **Table 2**). Both HFE (Model 1, onset: 310 ms, 7 clusters, largest: 472–792 ms, *p* < .001, peak *r* = 0.57) and MFCC (Model 2, onset: 310 ms, 3 clusters, largest: 442–998 ms, *p* < .001, peak *r* = 0.58) remained significant after controlling for F0. When mutually controlled, HFE (Model 3) showed substantially reduced clusters from 350 ms (3 clusters, largest: 578–628 ms, *p* < .001, peak *r* = 0.50), whereas MFCC (Model 4) remained robust with 10 significant clusters from 228 ms (*p* < .001, peak *r* = 0.56 at 458 ms), closely following the MVPA source decoding onset window (134–176 ms).

**Figure 4.**
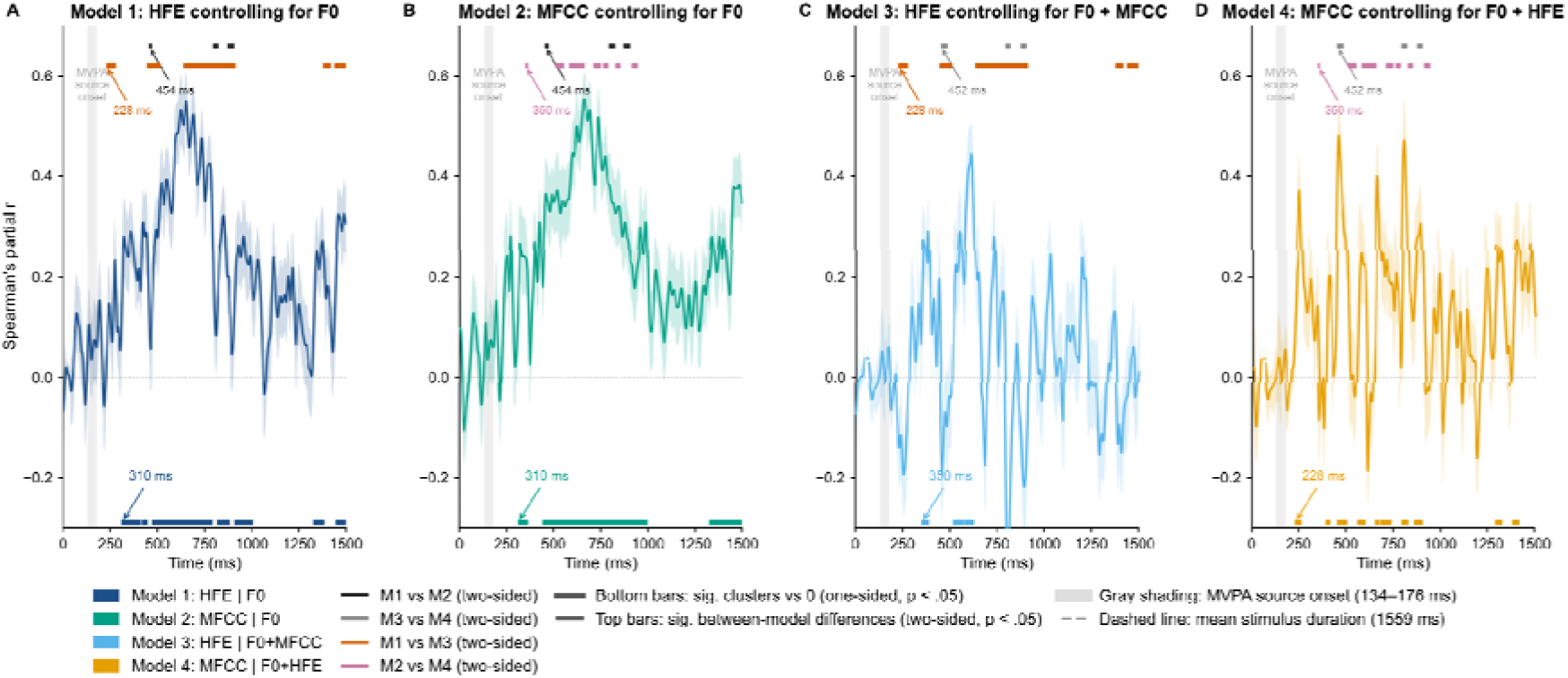
Partial RSA: Independent Acoustic Contributions to Neural Voice Source Structure. Partial Spearman correlations between neural and acoustic RDMs (*N* = 40, 2 ms resolution, 0–1500 ms). Each panel shows mean partial *r* (±1 SEM, shaded) after controlling for the specified features. Bottom colored bars: significant clusters vs. 0 (one-sided cluster permutation, *p* < .05); onset latencies annotated with downward arrows. Top bars: significant between-model differences (two-sided, *p* < .05); onset latencies annotated with upward arrows. Gray shading: MVPA source decoding onset (134–176 ms). Dashed line: mean stimulus duration (1559 ms). HFE = high-frequency energy (>4 kHz); MFCC = mel-frequency cepstral coefficients.

**Table 2.** Significant Clusters from Partial RSA: Independent Acoustic Contributions to Neural Voice Source Structure.

| Model | Feature | Controlling for | Cluster | Onset (ms) | Offset (ms) | Duration (ms) | $p$ | Peak $r$ | Peak time (ms) |
| --- | --- | --- | --- | --- | --- | --- | --- | --- | --- |
| Model 1 | HFE | F0 | 1 | 310 | 414 | 104 | 0.00 | 0.34 | 402 |
| Model 1 | HFE | F0 | 2 | 418 | 448 | 30 | 0.04 | 0.37 | 420 |
| Model 1 | HFE | F0 | 3 | 472 | 792 | 320 | < .001 | 0.57 | 648 |
| Model 1 | HFE | F0 | 4 | 814 | 882 | 68 | 0.01 | 0.34 | 854 |
| Model 1 | HFE | F0 | 5 | 908 | 1008 | 100 | 0.01 | 0.31 | 958 |
| Model 1 | HFE | F0 | 6 | 1326 | 1386 | 60 | 0.01 | 0.31 | 1380 |
| Model 1 | HFE | F0 | 7 | 1444 | 1500 | 56 | 0.01 | 0.36 | 1454 |
| Model 2 | MFCC | F0 | 1 | 310 | 364 | 54 | 0.03 | 0.34 | 324 |
| Model 2 | MFCC | F0 | 2 | 442 | 998 | 556 | < .001 | 0.58 | 662 |
| Model 2 | MFCC | F0 | 3 | 1326 | 1500 | 174 | < .001 | 0.42 | 1454 |
| Model 3 | HFE | F0 + MFCC | 1 | 350 | 390 | 40 | 0.00 | 0.46 | 352 |
| Model 3 | HFE | F0 + MFCC | 2 | 518 | 574 | 56 | 0.01 | 0.35 | 558 |
| Model 3 | HFE | F0 + MFCC | 3 | 578 | 628 | 50 | < .001 | 0.5 | 618 |
| Model 4 | MFCC | F0 + HFE | 1 | 228 | 266 | 38 | 0.01 | 0.45 | 256 |
| Model 4 | MFCC | F0 + HFE | 2 | 392 | 420 | 28 | 0.04 | 0.28 | 396 |
| Model 4 | MFCC | F0 + HFE | 3 | 452 | 508 | 56 | < .001 | 0.56 | 458 |
| Model 4 | MFCC | F0 + HFE | 4 | 560 | 606 | 46 | 0.02 | 0.3 | 576 |
| Model 4 | MFCC | F0 + HFE | 5 | 654 | 678 | 24 | 0.02 | 0.44 | 664 |
| Model 4 | MFCC | F0 + HFE | 6 | 682 | 742 | 60 | 0.01 | 0.3 | 708 |
| Model 4 | MFCC | F0 + HFE | 7 | 794 | 830 | 36 | 0.00 | 0.5 | 812 |
| Model 4 | MFCC | F0 + HFE | 8 | 860 | 908 | 48 | 0.01 | 0.34 | 896 |
| Model 4 | MFCC | F0 + HFE | 9 | 1290 | 1330 | 40 | 0.01 | 0.35 | 1294 |
| Model 4 | MFCC | F0 + HFE | 10 | 1382 | 1422 | 40 | 0.02 | 0.3 | 1404 |

*Note.* One-sided cluster permutation tests (10,000 permutations, $Z > 1.645$ , $\alpha = .05$ ). Onset and Offset: cluster boundaries in ms post-stimulus onset. Peak $r$ : maximum partial Spearman correlation within cluster. Model 1: HFE controlling for F0. Model 2: MFCC controlling for F0. Model 3: HFE controlling for F0 and MFCC. Model 4: MFCC controlling for F0 and HFE. HFE = high-frequency energy (>4 kHz); MFCC = mel-frequency cepstral coefficients (13 coefficients). $p < .001$ reported as < .001.

Between-model comparisons further clarified their relative contributions: M1 vs. M2 diverged from 454 ms; M1 vs. M3 from 228 ms, reflecting HFE’s reduction when MFCC is additionally controlled; M2 vs. M4 from 390 ms, showing comparatively less reduction for MFCC; and M3 vs. M4 from 452 ms, confirming MFCC’s stronger independent predictive power throughout the analysis window (**Figure 4**).

As such, despite its visual salience, HFE’s predictive power over neural human–AI discrimination is largely subsumed by MFCC, which emerges as the more robust and temporally earlier acoustic driver.

## 4. Discussion

The present study examined whether neural discrimination of human from AI voices depends on prosodic processing by comparing the temporal dynamics of voice source and prosody decoding. Three findings converged to suggest that prosody is not the primary basis of neural human–AI voice discrimination: voice source discrimination emerged substantially earlier than prosody decoding; prosodic information became neurally decodable only near utterance completion; and spectral envelope features rather than high-frequency energy drove early neural human–AI discrimination.

### 4.1. Neural Voice Source Discrimination Precedes Prosody Decoding, Supporting Retrospective Attribution

Our results support the retrospective attribution hypothesis: neural discrimination of human vs. AI voice source emerged rapidly (134–176 ms) and substantially preceded prosody decoding (1366–2066 ms), indicating that the brain distinguishes AI from human voices before prosodic information becomes available. This temporal dissociation offers a neural-level explanation for why listeners in behavioral studies report relying on prosodic cues when detecting audio deepfakes (Kühne et al., 2020; Rodero, 2017; San Segundo et al., 2025).

The early onset of voice source discrimination is comparable to the neural time course of accent perception (∼200 ms) (Goslin et al., 2012; Jiang et al., 2020; Llanos et al., 2021; Llanos et al., 2025; Romero-Rivas et al., 2015), which can be theorized within the perceptual magnet framework (Kuhl et al., 2008): just as native phonetic categories attract perception toward prototypical acoustic patterns and render non-native sounds perceptually deviant, listeners may possess internalized acoustic templates (Andics et al., 2010; Lavan et al., 2019; Maguinness et al., 2018) for human vocal production that flag AI voices as categorically anomalous (Englert et al., 2016; Nussbaum et al., 2025; Pisoni et al., 1985) within the first moments of exposure. Our data further propose that this discrimination operates below the level of conscious awareness, given that voice source was never task-relevant and participants were unaware of its neural discriminability.

Consistent with this interpretation, prior behavioral studies in which participants were subsequently asked to explain their detection judgments may have elicited retrospective accounts that reflect post-hoc reasoning (Fallshore & Schooler, 1995; Nisbett & Wilson, 1977; van Gaal et al., 2012) rather than the actual perceptual basis of discrimination. Our data offer a neural-level explanation for such reports. Listeners may attribute their detection to salient, consciously accessible features such as intonation or expressiveness (Kühne et al., 2020; Rodero, 2017; San Segundo et al., 2025), because rapid, pre-attentive voice source discrimination leaves no conscious trace. This prompts listeners to seek an explanation in prosodic features that are perceptually salient but neurally discriminable only near utterance completion.

### 4.2. Word-Level Prosodic Prototypes Generalize to the Utterance Level

As outlined in the Introduction, a central open question has been whether word-level prosodic prototypes generalize to sentence-level speech. Reverse-correlation work has identified distinct prototypes for dominance and trustworthiness from single-word utterances by deriving the mental representations driving listeners’ judgments of randomly perturbed pitch contours (Ponsot et al., 2018a), yet Ponsot et al. (2018b) acknowledged that generalization to longer utterances remains to be established. The present study provides convergent acoustic and neural evidence that such generalization is possible, at least for epistemic stance contrasts of confident vs. doubtful prosody, where confident speech corresponds to the certainty prototype identified at the word level (Goupil et al., 2021).

Using the same reverse-correlation approach, certainty and honesty were found to share a common prosodic signature extracted automatically and independently of listeners’ conceptual knowledge (Goupil et al., 2021). Confident prosody, as used here, reflects a similar epistemic state, suggesting a correspondence between our conditions and this word-level certainty prototype. Prior acoustic analyses of English confident and doubtful speech identified mean F0 of the final word as a critical distinguishing cue (Jiang & Pell, 2017), and the present Mandarin neural data converge on the same conclusion: prosodic information is encoded in and decoded from the terminal portion of the utterance. This provides evidence that the certainty-related prosodic prototype generalizes to the utterance level, with listeners integrating prosodic information over the full duration before accessing its social meaning. Whether prototypes for other social dimensions generalize similarly remains to be established (Knight et al., 2018; Ponsot et al., 2018b).

### 4.3. The Acoustic Basis of Human–AI Neural Discrimination Remains Elusive

Computational studies comparing human and AI-generated speech have identified a range of acoustic differences, including spectral envelope characteristics, jitter, shimmer, fundamental frequency variation, and pause patterns (Kulangareth et al., 2024; Nuska & Firdhous, 2025; Paul et al., 2017; Unoki et al., 2024), yet no single feature has emerged as a universal marker reliably distinguishing human from AI voices. Our mel spectrograms reveal that human voices carry more diffuse high-frequency energy than AI voices, suggesting high-frequency energy as a potential acoustic driver of neural discrimination. However, combining acoustic and neural RSA, we found that high-frequency energy was not the primary driver of neural discrimination. Instead, the broader spectral envelope captured by MFCC more robustly predicted neural human–AI structure. The earliest independent contribution of MFCC (228 ms) closely followed the MVPA source decoding onset (134–176 ms). This temporal convergence points to distributed spectral envelope features, rather than high-frequency energy, as a more plausible perceptual signal underlying early neural human–AI discrimination, consistent with computational studies identifying MFCC-based spectral envelope features as effective markers of synthetic speech (Nuska & Firdhous, 2025; Paul et al., 2017; Unoki et al., 2024).

Nevertheless, MFCC captures a broad composite of spectral envelope properties, including formant structure, spectral tilt, and coarticulation patterns arising from natural vocal tract dynamics (Abdul & Al-Talabani, 2022; Darch et al., 2008; Iskarous & Vietti, 2025), making it difficult to pinpoint which specific components carry the most diagnostic information for human–AI discrimination.

Future studies could address causality through acoustic resynthesis paradigms in which specific spectral dimensions are selectively manipulated or equalized across human and AI voices (Kitamura & Akagi, 1995; Koelewijn et al., 2023; Sell et al., 2015), directly testing whether individual acoustic features are necessary for neural and perceptual source discrimination. Stimulus-level RSA correlating trial-by-trial neural and acoustic dissimilarity across individual utterances could further identify moment-to-moment acoustic predictors of neural response (Lavan et al., 2024).

### 4.4. Limitations and Future Directions

The present study examined Mandarin Chinese speech, where monosyllabic structure allowed systematic control over utterance length and linguistic content. Whether the findings generalize to other languages with more variable syllable structure remains to be established. Future research examining listeners with no knowledge of the target language would provide a particularly strong test: if early neural human–AI discrimination persists without any linguistic familiarity, this would support a language-independent acoustic basis of voice source detection (Fleming et al., 2014; Perrachione et al., 2011).

A highlight of the present study is the use of voice cloning to match AI voices to individual human speaker identities, controlling for speaker-specific acoustic confounds, with stimuli drawn from a commercially available voice assistant service producing voices perceptually identifiable as AI-generated (Chen et al., 2026). One limitation is that the confident vs. doubtful prosodic manipulation does not fully correspond to the prosodic features listeners typically report when evaluating AI voices, such as monotonicity or reduced expressiveness (Kühne et al., 2020; Rodero, 2017; San Segundo et al., 2025), and the late-emerging decoding pattern observed here should not be generalized to all prosodic distinctions at the sentence level.

The fixed block order (human blocks 1–4, AI blocks 5–8) was motivated by the need to establish reliable human speaker identity representations as a baseline before introducing AI voices, as discussed in detail in Chen et al. (2025b). Although earlier blocks typically confer identity encoding advantages (Lavan et al., 2019; Xu & Armony, 2021; Zäske et al., 2014), our Checking phase accuracy confirmed that AI voices showed systematically lower recognition accuracy even when appearing in later blocks with cumulative learning advantages. This pattern could reflect inherent difficulty in encoding AI speaker identities, or alternatively, greater acoustic similarity among AI voices from the same synthesis system reducing within-category discriminability. The present study used the speaker identity learning task as a background context to examine implicit neural discrimination of voice source. Future studies could employ intermixed human and AI voice blocks with shallower tasks such as speaker sex judgments (Xu & Armony, 2021) to examine whether early neural discrimination of voice source is maintained under reduced attentional demands and without blocked exposure to each voice type.

Finally, the present sample comprised typical university students, and whether early neural discrimination of voice source generalizes to other populations remains to be examined, including those showing varied sensitivity to human vs. artificial voices such as older adults (Herrmann, 2023) and individuals with autism spectrum disorder (Kuriki et al., 2016).

## Acknowledgments

This research was supported by the National Natural Science Foundation of China (Grant No. 32471109), awarded to X. Jiang. The PhD studentship of W. Chen was supported by a McGill-CSC (China Scholarship Council) Joint Scholarship, part of which is sourced from the Natural Sciences and Engineering Research Council of Canada (NSERC) Discovery Grant (RGPIN-2022-04363) awarded to M. D. Pell.

## CRediT

W. Chen: Conceptualization, Data curation, Formal analysis, Investigation, Methodology, Visualization, Writing – original draft. M. D. Pell: Resources, Supervision, Funding acquisition, Writing – review and editing. X. Jiang: Conceptualization, Funding acquisition, Investigation, Project administration, Supervision, Writing – review and editing.

